# Temporal preparation and short-term temporal memory in depression

**DOI:** 10.1101/708370

**Authors:** Tzu-Yu Hsu, Hsin-Chien Lee, Timothy Joseph Lane, Marcus Missal

**Affiliations:** Graduate Institute of Mind Brain and Consciousness, Taipei Medical University, Taipei, Taiwan; Brain and Consciousness Research Center, TMU-Shuang Ho Hospital, New Taipei City, Taiwan; Department of Psychiatry, School of Medicine, College of Medicine, Taipei Medical University, Taipei, Taiwan; Department of Psychiatry, Shuang-Ho Hospital, New Taipei City, Taiwan; Research Center of Sleep Medicine, College of Medicine, Taipei Medical University, Taipei, Taiwan; Graduate Institute of Humanities in Medicine, Taipei Medical University, Taipei, Taiwan; Institute of Neurosciences (IONS), Cognition and System (COSY), Université catholique de Louvain (UCLouvain), Brussels, Belgium

## Abstract

Patient suffering of Major Depressive Disorder (MDD) often complain that subjective time seems to ‘drag’ with respect to physical time. This may point towards a generalized dysfunction of temporal processing in MDD. In the present study, we investigated temporal preparation in MDD. “Temporal preparation” refers to an increased readiness to act before an expected event; consequently, reaction time should be reduced. MDD patients and age-matched controls were required to make a saccadic eye movement between a central and an eccentric visual target after a variable duration preparatory period. We found that MDD patients produced a larger number of premature saccades, saccades initiated prior to the appearance of the expected stimulus. These saccades were not temporally controlled; instead, they seemed to reflect increased oculomotor impulsivity. In contrast, the latency of visually-guided saccades was strongly influenced by temporal preparation in controls; significantly less so, in MDD patients. This observed reduced temporal preparation in MDD was associated with a faster decay of short-term temporal memory. Moreover, in patients producing a lot of premature responses, temporal preparation to early imperative stimuli was increased. A reduction in premature saccades, however, was associated with reduced temporal preparation to late imperative stimuli.

In conclusion, reduced temporal preparation and short-term temporal memory in the oculomotor domain supports the hypothesis that temporal processing was altered in MDD patients. These observed deficits could reflect other underlying aspects of abnormal time experience in MDD.

## Introduction

Major Depressive Disorder (MDD) is often associated with an altered awareness of the passage time. Indeed, MDD patients often complain that subjective time is going by at a reduced pace compared with physical time (Gallagher, 2012; Ratcliffe, 2012; Vogel et al. 2018). This perturbed time awareness has led to a systematic investigation of time ‘perception’ using quantitative methods requiring an *explicit* judgment about durations. An explicit judgment about duration is the outcome of experimental tasks requiring comparison of time intervals, production or reproduction of a standard duration or verbal estimation (Msetfi et al., 2012; Droit-Volet, 2013; Davalos et al. 2018). This approach has led to conflicting results and the precise influence of depression on time perception remains elusive (see review in Oberfeld et al., 2014; Thönes and Oberfeld, 2015). However, temporal cognition is not limited to *explicit* temporal judgements. *Implicit* timing refers to the capacity to time actions based on temporal regularities in the environment (Coull and Nobre, 2008; Coull and Droit-Volet, 2018). It emerges in non-temporal tasks where temporal information is, nevertheless, essential to achieve optimal performance, as when making a saccade to a visual target. This implicit influence of elapsed time on movement preparation is often referred to as ‘temporal preparation’.

Temporal preparation is studied, classically, by using a warning stimulus (S_1_) that predicts the occurrence of an imperative stimulus (S_2_; Woodrow, 1914). The period between S_1_ offset and S_2_ onset is referred to as the foreperiod (‘FP’; Niemi 1981; Niemi and Näätänen, 1981). Temporal preparation builds-up while waiting during the FP and causes a shorter reaction time after S_2_ appearance.

Foreperiod duration could either remain constant, making the timing of S_2_ entirely predictable, or FP could vary randomly between different values drawn from a given probability distribution. If FP duration is randomly drawn from a uniform probability distribution, the latency of the motor response to S_2_ decreases with elapsed time. This ‘foreperiod effect’ is the behavioral measure of temporal preparation. Temporal preparation could be explained by hypothesizing that subjects estimate the hazard rate of the target defined as the probability that S_2_ will occur given that it has not occurred yet. As time elapses during the FP, the hazard rate of the S_2_ increases and sensorimotor systems could use that information to reduce reaction time (Trillenberg et al., 2000; Janssen and Shadlen 2005; Nobre et al. 2007). Temporal preparation could also be modulated by the previous FP’s experienced by the subject (Alegria and Delhaye-Rembaux, 1975; Los, 2010; Los and Van den Heuvel, 2001; Los et al. 2017). For instance, reaction time to S_2_ appearance during the current FP will tend to be shorter, if the previous FP was shorter. Therefore, short-term temporal memory (i.e. sequence effects) plays a crucial role in temporal preparation (Los et al. 2017). Accordingly, it has been shown that the FP effect on saccadic eye movements could be accounted for by the remaining trace of previous FP duration (Ameqrane et al. 2014). This influence of short-term memory on the RT-FP function could be altered given the known impact of depression on memory (Burt et al. 1995).

In the present study, we used saccadic eye movements to investigate temporal preparation. Indeed, precise and accurate control of the timing of eye movements is essential to ‘catch’ with the fovea the image of visual objects (Badler and Heinen 2006; de Hemptinne et al., 2007; Barnes, 2008; Collins and Barnes 2009). Furthermore, the FP effect is also observed using saccadic eye movements, if both S_1_ and S_2_ are visual targets and temporal preparation plays a major role in oculomotor control (Oswal et al. 2007; Ameqrane et al. 2014; Degos et al. 2018). Therefore, the aim of the present study was to examine the impact of depression on temporal preparation and short-term temporal memory using an oculomotor paradigm. Furthermore, we suggest how it could alter temporal cognition in MDD in general.

## Methods

### Ethical approval and informed consent

This study was approved by the joint institutional Review Board, Taipei Medical University, Taipei City, Taiwan (N201603080). Methods were carried out in accordance with relevant guidelines and regulations. All participants were informed about the purpose of the study and procedures before being asked to give informed consent. Written informed consent was obtained from all participants prior to their participation in this study.

### Experimental Design and Statistical Analysis

Subjects were facing an LCD screen which presented stimuli at a refresh rate of 60 Hz. An EyeLink 1000 infrared eye tracking system (SR Research, Mississauga, Ontario) was used to record eye movements at 1 KHz. Stimulus display and oculomotor data collection were synchronized on a frame-by-frame basis using Experimental Builder (SR Research, Mississauga, Ontario). Figure 1A depicts the time line of stimuli presentation on the screen facing the subject. Each trial started with an initial fixation period of a small empty box (1.4 × 1.4 deg) appearing on the screen for a random duration (850 ± 100 ms; Fig. 1A). At the end of this period four additional empty square ‘boxes’ appeared on the screen at an eccentricity of 8 deg. Afterwards, a green target (warning stimulus S_1_) was briefly presented in the central box for 50 ms. Extinction of the central target indicated to subjects the beginning of the foreperiod (FP). Subjects were required to fixate on the central box until another target was briefly and randomly presented for 50 ms, in one of the 4 eccentric boxes (imperative stimulus S_2_). One of 4 different FP durations (400, 900, 1400 and 1900 ms) was chosen randomly, each with the same probability (Probability=0.25). Subjects were required to wait until targets appeared in the eccentric box before making a saccade (black arrowhead on Fig. 1A). Saccadic latency (reaction time) was defined as the time elapsed between the appearance of the eccentric target and movement onset. Saccades that occurred before the appearance of the eccentric target (period indicated with a *red line* on Fig. 1A) are here referred to as ‘premature saccades’ (*red traces* on Fig. 1B). Saccades that occurred after the appearance of the eccentric target (period indicated with a *blue line* on Fig. 1A) are here referred to as ‘visually-guided saccades’ (*blue* traces on Fig. 1B). At the end of each trial, the display was replaced with a black screen and there was a variable inter-trial-interval of between 2000 and 2500 ms. A block contained 120 trials and each subject performed 2 blocks for a total of 240 trials. This paradigm is an oculomotor version of the well-known four choices serial reaction time task (i.e. 4-CSRTT) used in humans (Voon et al. 2014).

**Figure 1.**
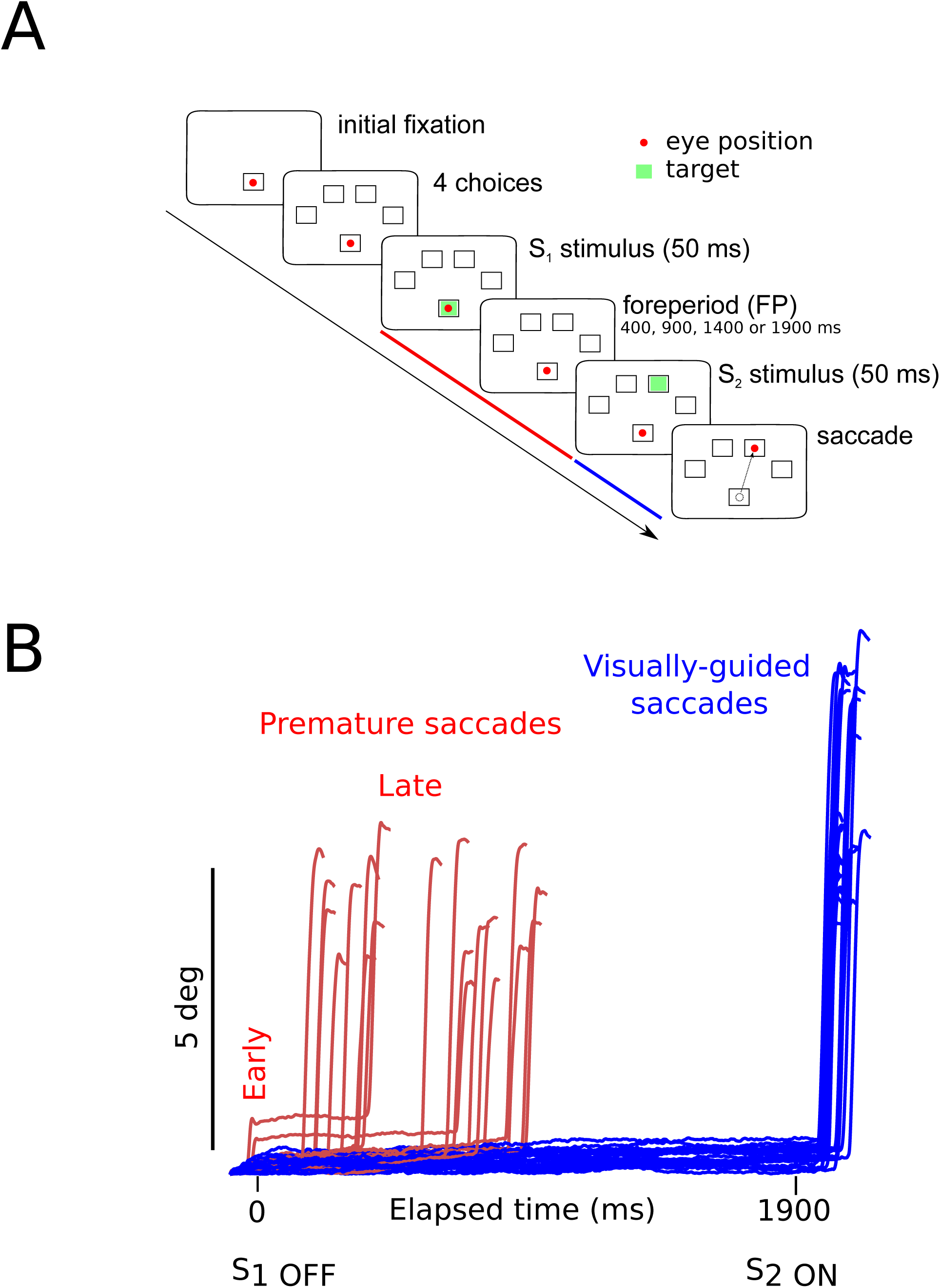
A: Oculomotor version of the 4-CSRTT task. Schematic representation of the visual display in front of subjects. Each trial started with the appearance of an empty box at the center of the screen. Subjects had to look at the center of the box (the *red disk* shows eye position) for a variable period of time. Thereafter, four additional empty ‘boxes’ (4 choices) appeared at an 8-deg eccentric position. After the appearance of the eccentric boxes, a warning stimulus S_1_ (*green* target) was flashed in the central box for 50 milliseconds. Extinction of the central target marked the beginning of the foreperiod (FP) that could last either 400, 900, 1400 or 1900 ms. At the end of the foreperiod, the imperative stimulus S_2_ appeared for 50 ms in one of the eccentric boxes randomly and the subject had to make a saccade (*dashed line*) to the cued eccentric box within a 400 ms grace period. B: Example of premature (*red traces*) and visually-guided (*blue traces*) saccades during a 1900-ms foreperiod in a control subject (subject #9). Time zero on the X-axis indicates extinction of the warning stimulus (S_1 OFF_). At the end of the FP, the imperative stimulus appeared (S_2 ON_). Note that most premature saccades tended to cluster during the early part of the FP. Small early premature saccades were sometimes followed by a later premature saccade. In this case, only the latency of the first premature saccade was measured. The period during which premature saccades were recorded is represented in *red* in Fig. 1A. The period during which visually-guided saccades were recorded is represented in *blue* in Fig. 1A.

The significance threshold α for all analysis was 0.05. All analyses were conducted using analysis of variance (ANOVA). Analyses were performed using SPSS 25 (SPSS Inc., Chicago, IL). Mixed-design ANOVA’s were used, unless otherwise specified. In mixed-design ANOVA’s, subject identity was used as a random factor to account for the influence of uncontrolled, between-subject variability. Results are presented as mean ± standard error of the mean, unless otherwise specified. Because saccadic reaction time distributions are often non-normal, we used a logarithmic transform of saccadic RT for statistics. For presentation purposes, however, untransformed saccadic latencies in milliseconds are presented on figures.

### Patients

Twenty-nine patients diagnosed with MDD (24 females; 38.4 ±2.5 years old, n=29) were recruited by the Department of Psychiatry at Taipei Medical University Shuang-Ho Hospital, located in New Taipei City, Taiwan. The MINI-international neuropsychiatric interview (Sheehan eat al., 1998), was used to confirm the diagnosis of current major depressive episode, to detect suicidal risk, and to exclude patients with psychotic symptoms and any comorbid mental disorder or substance use disorder according to the Diagnostic and Statistical Manual of Mental Disorders, Fifth Edition. Patients with poor visual acuity and comorbid medical conditions including neurological disorders (e.g. stroke, seizure, traumatic brain injury, post-brain surgery), brain implants (neurostimulators), cardiac pacemakers, or pregnant were also excluded. The Beck Depression Inventory (BDI-II; Beck et al. 1996), and the Generalized Anxiety Disorder 7 (GAD-7; Spitzer et al. 2006) were administered to evaluate the severity of depression and anxiety. BDI-II score for patients in this study was 30.1 ±13.4 and depressive symptoms duration was 9.2 ± 10.2 years. In addition, the Barratt Impulsiveness Scale 11 (BIS-11; Patton et al. 1995) was used to quantify Impulsive level on each individual. All patients but 7 were on medication at the time of testing (see Table 1).

**Table 1.** Summary of drug treatments received by patients. Each line represents one patient. Columns show treatments received by each patient (1 to 6).

| patient | 1 | 2 | 3 | 4 | 5 | 6 |
| --- | --- | --- | --- | --- | --- | --- |
| p01 | SNRI | Non-BDZ | BDZ | ATA | BDZ |  |
| p02 | MRA | Non-BDZ |  |  |  |  |
| p03 | SSRI | BDZ |  |  |  |  |
| p04 | BDZ | Non-BDZ | NDRI |  |  |  |
| p05 | **NIL** |  |  |  |  |  |
| p06 | ATA | SSRI |  |  |  |  |
| p07 | NDRI |  |  |  |  |  |
| p08 | MRA |  |  |  |  |  |
| p09 | SSRI | ATA |  |  |  |  |
| p10 | SSRI | ATA | NDRI |  |  |  |
| p11 | SNRI | ATA |  |  |  |  |
| p12 | SSRI |  |  |  |  |  |
| p13 | SNRI | Non-BDZ | BDZ |  |  |  |
| p14 | **NIL** |  |  |  |  |  |
| p15 | SSRI |  |  |  |  |  |
| p16 | **NIL** |  |  |  |  |  |
| p17 | **NIL** |  |  |  |  |  |
| p18 | SNRI | Non-BDZ | Non-BDZ | SARI | ATA | ATA |
| p19 | SSRI |  |  |  |  |  |
| p20 | SSRI | BDZ | BDZ |  |  |  |
| p21 | SNRI | ATA | BDZ | BDZ | SARI | BDZ |
| p22 | SSRI | BDZ | ATA |  |  |  |
| p23 | Non-BDZ | BDZ |  |  |  |  |
| p24 | NDRI |  |  |  |  |  |
| p25 | **NIL** |  |  |  |  |  |
| p26 | **NIL** |  |  |  |  |  |
| p27 | SNRI | BDZ | Non-BDZ | ATA |  |  |
| p28 | SNRI | ATA |  |  |  |  |
| p29 | **NIL** |  |  |  |  |  |

*ATA*, Atypical antipsychotic; *BDZ*, Benzodiazepine; *SNRI*, Serotonin-norepinephrine reuptake inhibitor; *MRA*, Melatonin receptor agonist; *NDRI*, Norepinephrine-dopamine reuptake inhibitor; ***NIL***, no pharmacological treatment; *Non-BDZ*, Non-benzodiazepine hypnotic; *SARI*, Serotonin antagonist and reuptake inhibitor; *SSRI*, Selective serotonin reuptake inhibitor;

### Controls

Twenty-nine healthy control participants (27 females, 37.7 ± 2.4 years old) without any current or history of neurological or psychiatric disorder, or use of psychotropic medication, were recruited from the community. They were matched for age and gender, except for two healthy control participants, whose gender did not match the patients.

## Results

### Demographics and clinical characteristics

Average age of control subjects was 37.7±2.4 (n=29) and 38.4±2.5 (n=29) for patients. Age did not differ between groups (F[1, 56]=0.048; *p*=0.827; n=29). Beck’s Depression Inventory-II (BDI-II, Beck et al. 1996) was used to assess the severity of depressive symptoms in healthy control subjects and patients. In controls, average scores were 7.6±1.4 (n=25/29 controls; 4 untested controls) and 30.1±13.4 in patients (n=22/29; seven untested patients). These group scores were statistically different (χ^2^=20.455; *p*<0.001). Trait impulsivity was estimated using the Barratt Impulsiveness Scale (BIS-11, Patton et al. 1995). In controls, average total scores were 64.6±1.2 (n=29/29 controls) and 71.3±2.0 in patients (n=27/29). These between-group scores differed significantly, (χ^2^=4.577; *p*=0.032) with patients scoring higher than controls.

### Saccades in controls and MDD patients

In the oculomotor version of the 4-CSRTT task that we used, subjects were instructed to keep looking at the central fixation box and wait for the appearance of the eccentric target to initiate a visually-guided saccade to one of four possible locations. Figure 1B shows raw eye movement traces recorded in one subject. Most saccades were visually-guided, as expected, in accord with the instructions provided to subjects (*blue traces* on Fig. 1B; 76% visually-guided saccades, 9451/12452 saccades all subjects included; average latency 239±1ms, n=5377 saccades; MDD: 259±1ms, n=4074 saccades). Average saccadic latency of visually-guided movements did not differ between groups (F[1, 55.154]=1.061; *p*=0.308). MDD patients produced normal latency saccades. We did observe, however, a large number of saccades initiated before the appearance of the target S_2_ that will be referred to as ‘premature’ saccades. Across all subjects, the percentage of premature responses reached 24%. But the percentage of premature saccades was more than twice as high in patients than in controls (in controls: 14±2 %, n=29 subjects, 937 premature saccades/6620 trials; in patients: 31±5 %, 2064 premature saccades/6776 trials) and there was a significant main effect of subject group on this percentage (F[1, 56]=8.4; *p*=0.005). In controls, the percentage of premature responses remained lower than 40% in most subjects, whereas the percentage of premature saccades reached as high as 80% in the most extreme cases of depressed patients. Figure 2 shows the latency distribution of premature saccades for all controls (Fig. 2A) and patients (Fig. 2B). The origin of the latency distribution axis is the extinction of the central target. It can be seen that the latency distribution presented two peaks: the first peak corresponds to very early premature responses, occurring near the time of fixation period offset. The second peak corresponds to later responses, likely triggered by S_2_ expectation. The observed distribution is bimodal; the data sample separates into early premature (response latency <100 ms) and late premature saccades (response latency > 100 ms). With respect with the total sample of premature responses (early + late responses), the percentage of early premature saccades was, on average, 17±3% in controls (n=21/29 subjects) and 13±2% in patients (n=24/29 subjects). Although MDD patients initiated more premature responses (see above), the percentage of early premature responses with respect to the total number of premature responses was the same between groups (F[1,43]=0.406; *p*=.527). In controls, the mean latency of early premature responses was 20±4 ms (n=21/29 subjects; 8 controls produced no premature responses); in patients, 17±3 ms (n=24/29 subjects; 5 patients produced no premature responses). Latency did not differ significantly between patients and controls (F[1, 43]=.880; *p*=.353).

**Figure 2.**
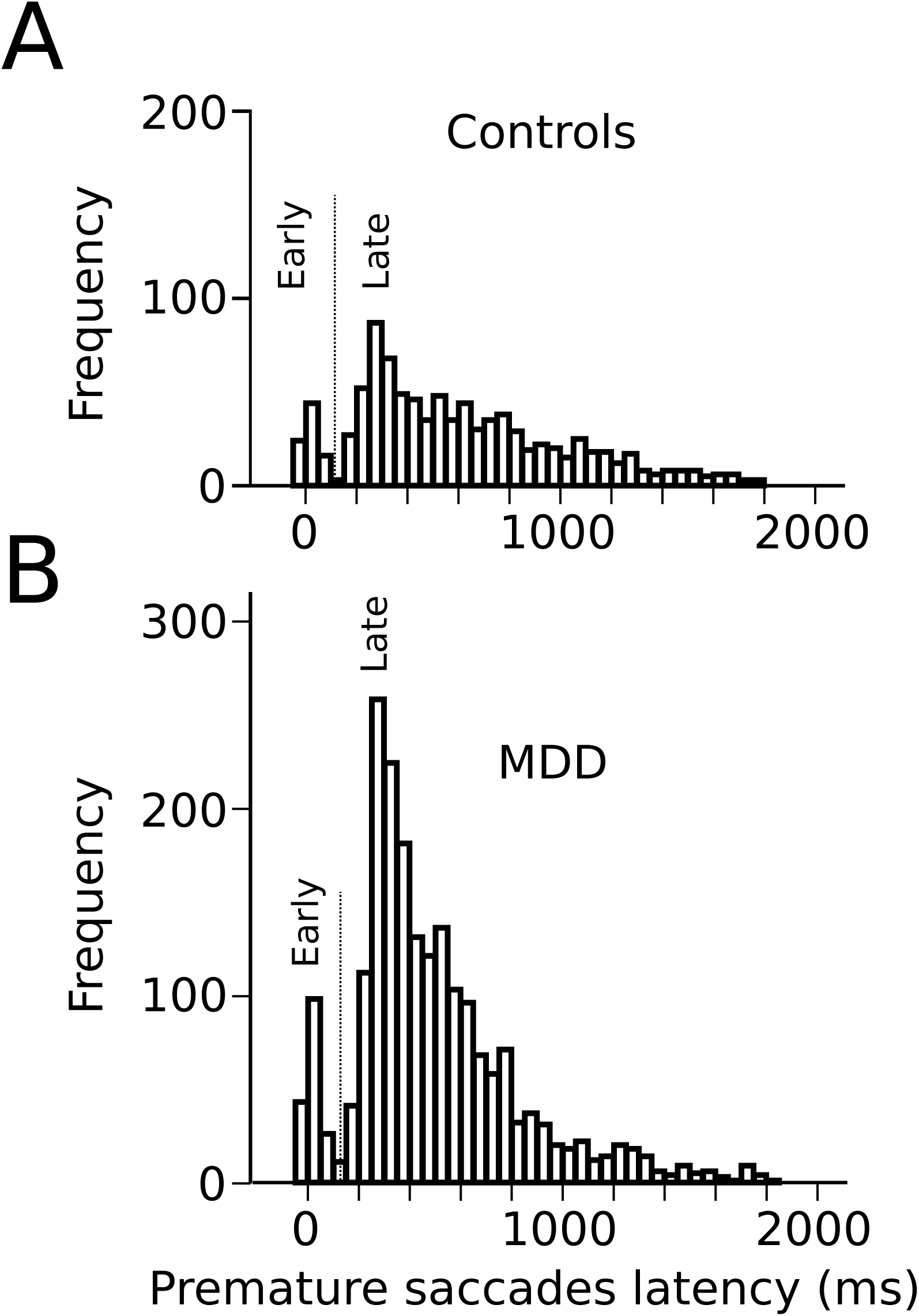
A: Latency distribution of all premature saccades in controls pooled together. B: Latency distribution of all premature saccades in MDD patients. Time zero on the X-axis represents disappearance of the S_1_ stimulus. The *vertical dashed line* indicates the threshold for separating early and late premature responses.

Late premature saccades were planned after the extinction of the central fixation target, during expectation of the eccentric target. These late responses were more frequent than early responses and were not uniformly distributed: i.e. they did not occur with the same probability during the FP. Indeed, most observations were clustered between 200 and 400 ms after S_1_ offset. This observation suggests that they were triggered by the expectation of S_2_ appearance after a short FP (400 ms). The percentage of late premature saccades with respect to total premature responses was, on average, 88±3% in controls (n=29 subjects) and 90±2% in patients (n=29 subjects). The percentage of late premature saccades did not differ between groups (F[1, 56]=0.386; *p*=.537). In controls, late premature saccade latency was, on average, 667±27 ms (n=29 subjects) and 598±24 ms in patients (n=29 subjects). However, no between-group statistical difference was observed (F[1, 56]=3.656; *p*=.061).

In summary, visually-guided saccadic latency was similar for both groups. We observed, however, more than twice the percentage of premature saccades in patients than in controls. Although the BIS-11 score was on average higher in MDD patients, we found no correlation with the percentage of premature saccades. This result suggests that the BIS-11 score and the percentage of premature saccades are biomarkers of two different facets of impulsivity. Therefore, in the present study ‘trait impulsivity’ will be used to refer to what is measured using questionnaires like the BIS-11. ‘Oculomotor impulsivity’ will be used to refer to an increased propensity to generate premature responses.

### Temporal preparation in health and depression

Temporal preparation causes a regular decrease of saccadic latency with increasing FP duration (Ameqrane et al. 2014), similar to what is found in other sensorimotor modalities (Los et al. 2014). The resulting curve will be referred to as the ‘RT-FP function’. Figure 3A shows the RT-FP functions for both groups. It can be observed that saccadic latencies were similar, if current FP duration was 400 ms. But there was an increasing difference between groups for FP duration longer than 400 ms. We found a statistically significant main effect of FP duration (F[3, 9394.324]=274.374; *p*<0.001) on saccadic latency, but no significant main effect on average saccadic latency (F[1, 55.082]=0.472; *p*=0.495). The interaction between FP duration and subject group was significant (F[3, 9394.324]=22.715; *p*<0.001). Importantly, these results show that *average* reaction time was statistically similar in controls and MDD subjects, but that the FP effect was significantly reduced in MDD patients. This result suggests reduced temporal preparation in MDD.

**Figure 3.**
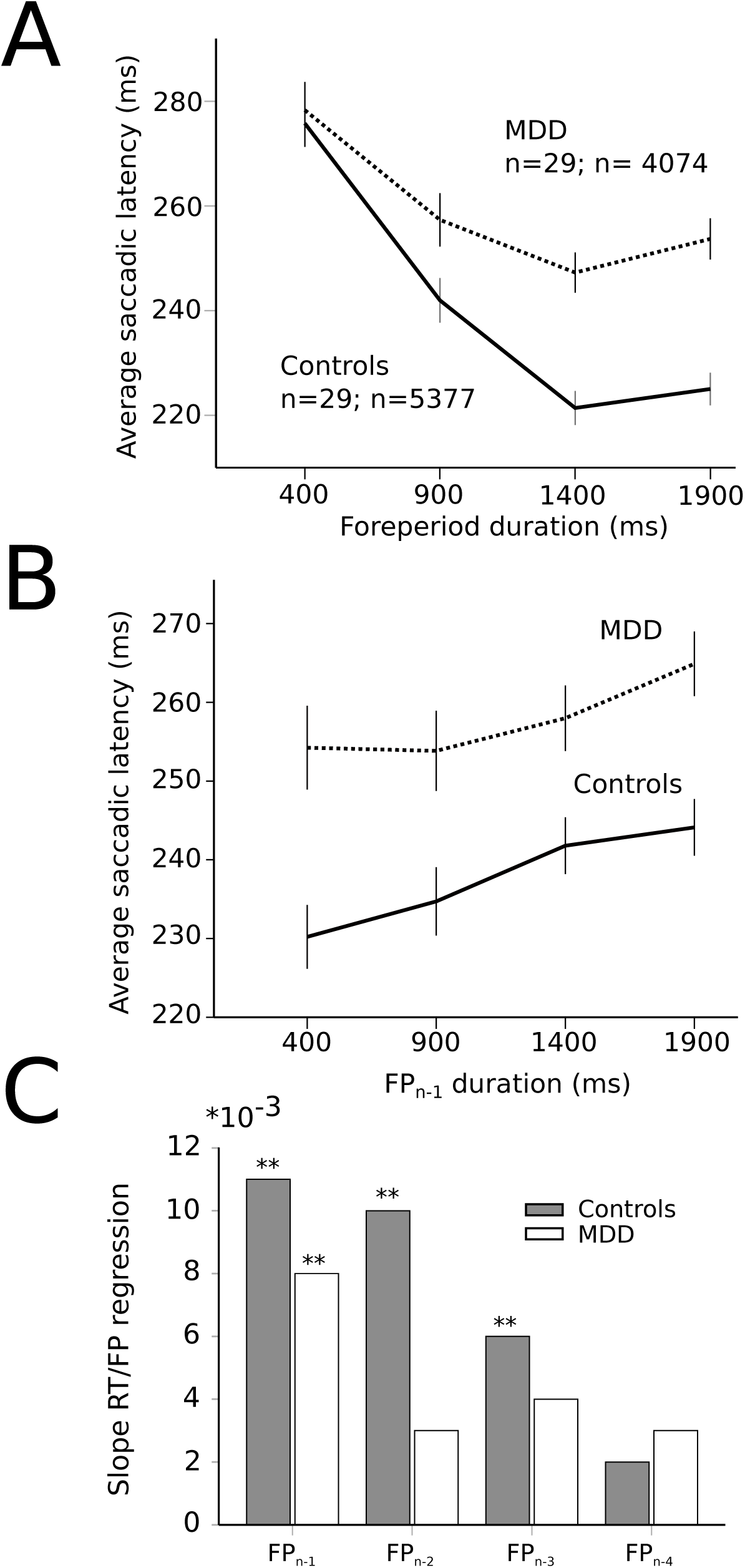
A: RT-FP function in controls (*continuous* line; n=29 subjects; n=5377 saccades) and MDD patients (*dotted* line; n=29 subjects; n=4074 saccades). Average values of saccadic latency (in milliseconds) and 95% confidence interval. Note the shallower slope of the function between 400 and 1400 ms FP durations. B: Relationship between previous foreperiod duration (FP_n-1_) and saccadic latency in controls (*continuous* line) and MDD patients (*dotted* line). Same data set as on Fig. A. C: Graphical representation of the slope of the linear relationships between previous FP’s and saccadic latency for an increasing number of trials back into the past (1 to 4) in controls (*dark* bars) and MDD patients (*open* bars). Same data set as on Fig. A.

In order to better understand the origin of the different RT-FP functions in MDD and controls, we analyzed the influence of previous FP duration (FP_n-1_) on saccadic latency during the current FP. For instance, if current FP duration was 400 ms and FP_n-1_ was 400 ms as well, saccadic latency could be shorter than if the same FP was preceded by FP_n-1_ of 1900 ms (see Fig. 3B). Statistically, we found that there was a significant main effect of FP_n-1_ duration (F[3, 9314.030]=22.841; *p*<0.001) on saccadic latency, but not subject group (F[1, 55.229]=1.077; *p*=0.304). The interaction between FP_n-1_ duration and subject group was not significant ([F(3, 9314.030)=1.633; *p*=0.179]). These results suggest that the FP_n-1_ effect was present and similar in both controls and patients. Therefore, the reduced FP effect that we found could not be due to a reduced influence of FP_n-1_ duration held in short-term memory. More than one step back into the past, however, could still play a significant role in the timing of eye movements. Therefore, we investigated whether FP_n-1_, FP_n-2_, FP_n-3_, FP_n-4_ could also play a significant role in determining saccadic latency during the current FP_n_ in both groups. We found that FP_n-1_, FP_n-2_, FP_n-3_ and FP_n-4_ played a significant role in determining saccadic latency in controls (FP_n-1_, F[3, 4935.335]=16.090; *p*<0.001; FP_n-2_, F[3, 4935.316]=9.382; *p*<0.001; FP_n-3_, F[3, 4935.283]=4.126; p=0.006; FP_n-4_, F[3, 4935.237]=3.562; *p*=0.014). But only FP_n-1_ played a significant role on saccadic latency in MDD patients (FP_n-1_, F[3, 3654.868]=6.270; *p*<0.001). Therefore, the reduced FP effect found in patients could be attributed to a different processing of past FP durations, with a faster decay of the memory trace of previous FP’s in MDD patients. This is further illustrated on Fig. 3C, which shows the value of the slope of a multiple linear regression analysis between previous FP’s and saccadic latency. Star symbols indicate when slopes were significantly different from zero. This figure shows a progressive extinction of the influence of previous FP’s on movement latency in MDD patients.

Analyses presented above suggest that short-term temporal memory of previous FP’s affected saccadic latency in MDD patients differently than in controls. Temporal preparation, however, could also be guided by a sense of elapsed time during the current FP. Therefore, we applied a hierarchical linear regression analyses in both groups using two different models. In controls, the first model contained FP_n-1_, FP_n-2_, FP_n-3_ and FP_n-4_ as independent factors. The second model contained FP_n-1_, FP_n-2_, FP_n-3_, FP_n-4_ and FP_n_ as independent factors and will be referred to as ‘model 2’. The aim was to determine the contribution of both short-term temporal memory (FP_n-x_) and current foreperiod (FP_n_) in movement latency. In controls, we found that both model 1 (F[4, 5213]=17.269; *p*<0.001) and model 2 (F[5, 5212]=89.276; p<0.001) provided a significant explanation of observed variance. The coefficient of determination for model 1 was r^2^ = 0.013 (*p*<0.001); however, the coefficient of determination for model 2 was higher with r^2^ = 0.079. Therefore, r^2^ variation due to adding current FP_n_ in the model was approximately 7% (0.066) and the F-value variation related to this addition was significant (F[1, 5212]=372.384; *p*<0.001). In patients, we found that both model 1 (F[1, 4038]=12.559; *p*<0.001) and model 2 (F[2, 4037]=38.633; *p*<0.001) also provided a significant explanation of the variance observed.

The coefficient of determination for model 1 was weak r^2^ = 0.003 but significant (*p*<0.001). But the coefficient of determination for model 2 was higher with r^2^ = 0.018 and the r^2^ variation due to adding FP_n_ in the model was approximately 2% (0.016). F-value variation related to the addition of FP_n_ in the model was significant (F[1, 4037]=64.511; *p*<0.001). In summary, the FP effect was observed in both groups but in MDD patients memory of previous FP’s declined faster, given that only FP_n-1_ played a significant role on saccadic latency. Using a hierarchical model analysis, we showed that the model including previous FP’s and current FP_n_ explained a larger proportion of the variance of saccadic latency. Influence of the current FP was present in both controls and patients but its influence was weaker in the latter group. Therefore, the reduced influence of current FP on saccadic latency co-occurred with a reduced short-term temporal memory in MDD patients.

### Was short-term memory in the spatial domain similarly affected in MDD patients?

If the answer to this question were positive, then observed effects were not specific to temporal preparation. Instead, they might reflect a more general short-term memory impairment in patients. In order to determine whether there was a significant effect of previous target *position*, we compared saccadic latency when the target appeared at the same or at a different spatial location during the previous trial. We hypothesized that there could be a response facilitation by repetition in the temporal domain only, but not in the spatial domain. Indeed, we found no evidence of a significant interaction effect between subject group and previous target *location* on saccadic latency (F[1, 9318.688]=0.081; *p*=0.776). We applied the same approach to temporal preparation by comparing saccadic reaction time when FP_n_ and FP_n-1_ were the same or different. In the temporal domain, there was a significant interaction between subject group and FP duration on saccadic latency (same or different; F[1, 8907.674]=4.077; *p* =0.043). These results show that *temporal* short-term memory was selectively affected in patients, but the same was not true for *spatial* short-term memory of target location.

### Influence of oculomotor impulsivity on temporal preparation

The shape of the RT-FP function could be influenced by both oculomotor impulsivity and depression. Therefore, we compared the RT-FP functions in low and high oculomotor impulsivity patients and controls. Patients were categorized into 2 groups, using the median of the percentage of premature saccades distribution in this group (median = 21%). The low impulsivity MDD group produced less than 21% premature responses (n=15 subjects; n=2864 saccades); the high impulsivity MDD group, more than 21% (n=14 subjects; n=1210 saccades). We re-examined the FP effect within these 2 groups. Figure 4A shows that the asymmetric FP effect was present in the low impulsivity group, but it was considerably altered in the high impulsivity group. Accordingly, a significant interaction between impulsivity group and FP duration on saccadic latency was found (F[3, 4048.361]=4.655; *p* =0.003). In high oculomotor impulsivity patients, saccadic latencies for the 400 ms duration FP were reduced by approximately 30 ms. Figure 4B shows the same analysis applied to control subjects (median percentage of premature saccades: median = 10%). The RT-FP function in high (n=15 subjects; n=2313 saccades) and low impulsivity (n=14 subjects; n=3064 saccades) control subjects also significantly differed (significant interaction between impulsivity group and FP duration on saccadic latency; F[3, 5342.181]=4.125; *p* =0.006), but to a lesser extent. Indeed, as already mentioned, general impulsivity was lower in this group; therefore, its influence on the RT-FP relationship was less. In summary, impulsivity reduced temporal preparation in both controls and patients, with a more pronounced effect in the latter group.

**Figure 4.**
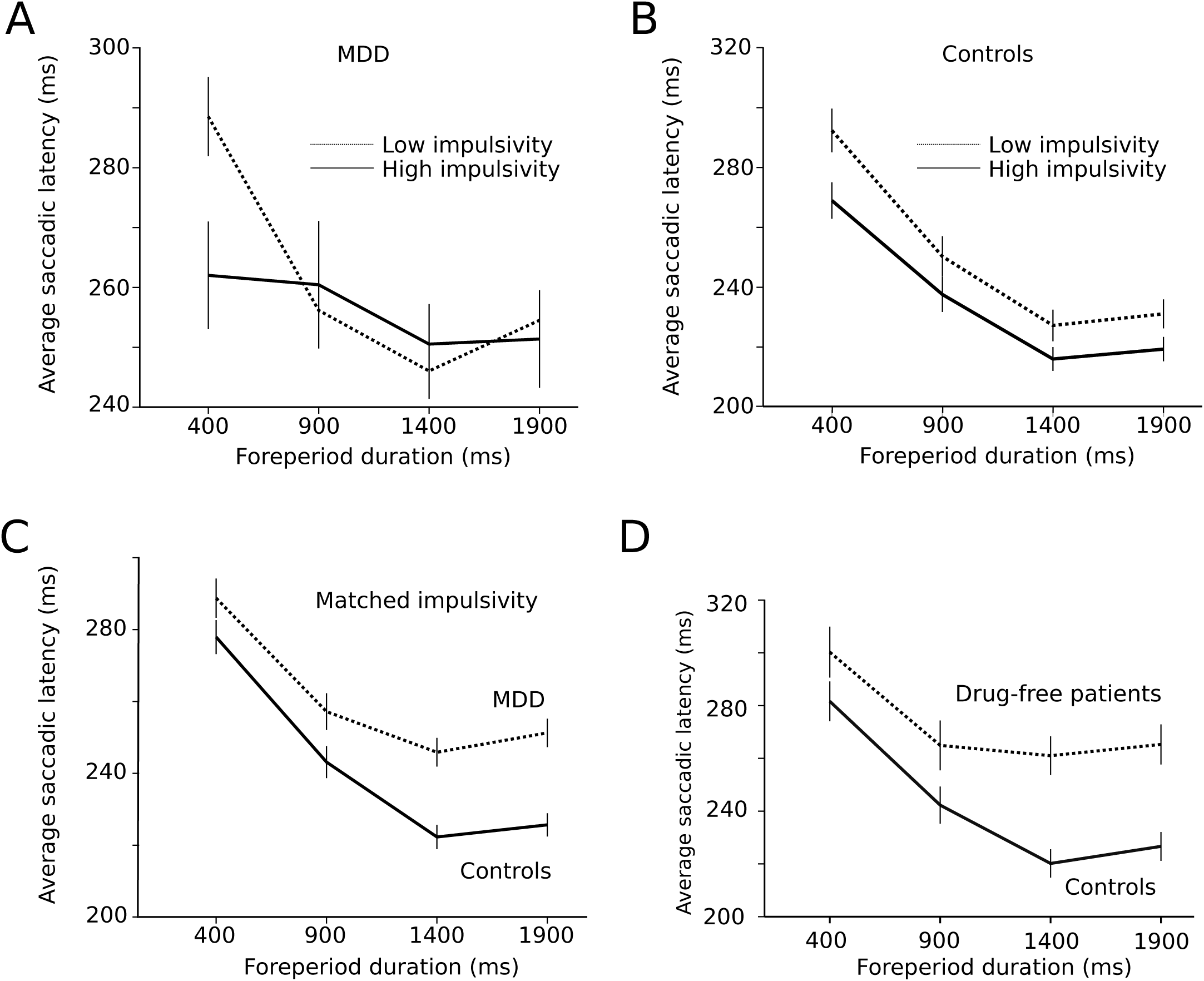
A: Influence of impulsivity on the RT-FP function in MDD patients (low impulsivity; n=15 subjects; n=2864 saccades; high impulsivity group: n=14 subjects, n=1210 saccades). B. Same relationship in controls (low impulsivity; n=15 subjects; n=2864 saccades; high impulsivity group: n=14 subjects, n=1210 saccades). C. RT-FP function for patients (n=19, 9 subjects removed; n=3546 saccades) and controls (n=26, 2 subjects removed; n=4972 saccades) that were matched for oculomotor impulsivity. **D:** RT-FP function in 7 untreated patients (n=1305 saccades) and 7 age-matched controls (n=1530 saccades).

The perception of elapsed time, however, could also be modified by depression, independent of impulsivity. In order to test this hypothesis, patients and controls were matched for oculomotor impulsivity. If the shape of the RT-FP relationship was different for the same oculomotor impulsivity, then depression is affecting temporal preparation. In order to obtain matched oculomotor impulsivity groups, we removed patients and subjects producing more than 30% premature responses and identified two overlapping groups— controls (n=26, 2 subjects removed; n=4972 saccades) and depressed patients (n=19, 9 subjects removed; n=3546 saccades) producing 12% and 14% of premature responses respectively. Figure 4C shows that the RT-FP functions were different, nevertheless, in patients and controls (significant interaction; F[3, 8467.292]=13.082; *p* <0.001) with a reduced FP effect in patients. Therefore, the shape of the RT-FP function could be altered by both oculomotor impulsivity and depression.

One additional factor that could influence temporal preparation is drug treatments administered to MDD patients. Most subjects (22/29) received several drug treatments that could have influenced temporal preparation (see Table 1). Due to ethical considerations, it would not have been acceptable to ask patients to suspend their treatment for research purposes. But 7 subjects were investigated before the start of the treatment and temporal preparation could be estimated (n=1305 saccades). Figure 4D shows a strong alteration of the RT-FP function for drug-free patients, compared to age-matched controls (n=1530 saccades). This subset of 7, untreated subjects presented a similar alteration of the RT-FP function as was found when all subjects were pooled together (see Fig. 3A), and a significant interaction between group and FP duration on saccade latency was found (F[3, 2815.141]=10.639; *p*<0.001). This result suggests that temporal preparation alteration in depression is a robust observation, and that it is not caused by drug treatment. We found no correlation between the BDI score and the slope of the RT-FP relationship (Pearson correlation, -0.032, *p*=0.88, n=22).

### Influence of trait impulsivity on temporal preparation

In the present study, the percentage of premature responses was used to evaluate oculomotor impulsivity. We found that this facet of impulsivity profoundly influences temporal preparation. We also evaluated ‘trait’ impulsivity using the BIS-11 score (see above). Could the BIS-11 score predict the shape of the RT-FP function? In order to answer to this question, we divided MDD patients into a high and low impulsivity groups based on the median of the total BIS-11 score. The same ANOVA mixed-design procedure was applied to test whether the RT-FP function was different between groups. However, we found that there was no main effect of trait impulsivity on saccadic RT (F[1, 24.250]=0.001; p=0.973). Moreover, no significant interaction was found between trait impulsivity group and FP duration (F[3, 3777.549]=2.235; *p*=0.082). Therefore, temporal preparation was not altered by trait impulsivity.

## Discussion

The aim of this study was to determine how temporal preparation and short-term temporal memory were affected in MDD. In general, we found that temporal preparation was reduced in MDD compared with age-matched healthy controls. Moreover, we found that temporal preparation depended on both the duration of the current FP and short-term memory of previous FP’s. In MDD, the influence of current FP duration was reduced and the decay of short-term temporal memory was faster. Indeed, the influence of previous FP’s was 3-4 trials deep in controls, but only 1 trial deep in depressed patients. A significant reduction of the FP effect was found in 7 MDD patients who were not undergoing pharmacological treatment; this finding suggests that abnormal temporal preparation occurs independently of therapeutic drugs. In addition, we found that lack of inhibition also influenced temporal preparation. The RT-FP function was statistically flat in high oculomotor impulsivity patients, as evidenced by the strong reduction of saccadic latencies for the short 400 ms FP. On the contrary, in low oculomotor impulsivity patients the latency of saccades for the long FP’s was increased.

### The origin of temporal preparation

We suggest that psychomotor retardation could not fully explain observed results. Indeed, our analysis revealed that *average* reaction time to visual targets was not statistically different between groups. The multiple trace theory of temporal preparation (abbreviated as ‘MTP’; Los et al. 2017) suggests that the shape of the RT-FP function could be explained by a single learning mechanism based on previously experienced FP’s. For instance, if the current FP was preceded by a longer FP_n-1_ then reaction time could be increased. Across trials a memory trace builds-up that reflects the distribution of previous FP’s and the S_1_ stimulus triggers a recall of this composite memory trace during the current FP. Most recent traces play a larger role in preparation given that they are more readily available. Therefore, any reduction of the storage of previous FP’s should cause a change of the slope of RT-FP function, just as is observed in depressed patients in the present study. Moreover, we found that duration of the current FP was also significantly affecting the shape of the RT-FP function. Indeed, the hierarchical modeling analysis revealed that model 2 explained a larger proportion of the variance of RT than model 1 that did not include current FP. This observation suggests that the sense of elapsed time during the current FP could also be reduced by depression. Temporal preparation could have a dual origin (Vallesi and Shallice 2007; Vallesi et al. 2007a; Vallesi et al. 2007b) both affected in MDD.

Inhibitory control, likewise, plays also a crucial role while waiting for an expected event (Logan et al. 1997). Motor responses occurring before the appearance of the imperative S_2_ stimulus must be actively suppressed. If inhibition was not strong enough, then a premature response occurred. Premature responses could be driven by anticipation of S_2_ appearance. Indeed, the shaping of temporal preparation could require time-bound inhibitory functions (Burle et al. 2010; Correa et al. 2010; Los et al., 2014; Lebon et al. 2016). If this hypothesis is true, then the latency distribution of premature responses during the FP (the ‘mirror image’ of inhibition) should reveal an increased probability of premature responses at each likely timing of S_2_ appearance. Alternatively, the probability of occurrence of a premature response should be constant, if inhibition is uniform during the preparatory period (Greenhouse et al., 2015; Duque et al 2017). In contrast with these predictions, we observed that premature saccades clustered between 200 to 400 ms after S_1_ offset. This observation is not reconcilable with existing models. Accordingly, we suggest that two hypotheses could explain this observation. First, there could be an increased expectancy of S_2_ appearance after a short FP. This increased expectancy could be caused by a more salient representation of the short FP in memory. Second, response inhibition might be reduced at the beginning of the FP. But the observation that short-term temporal memory was reduced in MDD patients does not support the first hypothesis. Indeed, a reduced memory span and an increased saliency of memory traces of short FP’s are inconsistent. On the other hand, the second hypothesis is supported by observations made in humans and rodents. It has been suggested that the frontal and prefrontal cortices could exert inhibitory control over behavior, cognition and emotions although there is not yet a consensus about the networks involved in the different domains of inhibition (Jahanshahi and Rothwell 2017). Interestingly, Narayanan et al. (2006) have shown that in rat reversible inactivation of the dorsomedial prefrontal cortex caused an increase in premature responses and an accelerated reaction time after a short foreperiod, just as was observed in the present study. Therefore, premature saccades might reveal a lack of top-down inhibitory control exerted by the prefrontal cortex; this would be particularly important at the beginning of the FP. Later, during the FP, an excitatory drive could prepare movement initiation when the imperative stimulus occurs. The inhibitory control normally opposes this excitatory drive. But if the inhibitory drive were reduced then the excitatory drive could trigger either a premature response during the FP or a short latency saccade to the imperative visual target during a 400 ms FP. An increased occurrence of premature responses is often observed in psychiatric disorders and addictions where impulsivity is increased (Dalley and Robbins, 2017; Paasche et al. 2018). Among psychiatric diseases, MDD seems to be a special case. It is usually associated with heightened inhibitory control and reduced motor activity (American Psychiatric Association, 2013). But increased impulsivity has also been reported (Corruble et al. 2003) and top-down inhibitory control is likely to be affected by MDD (Palmwood et al. 2017). This hypothetical reduction of top-down inhibitory control in MDD could be related to reduced serotoninergic neurotransmission. Indeed, it has been shown in humans that a dietary tryptophan depletion procedure that reduces serotonin neurotransmission causes waiting impulsivity (Worbe et al. 2014; Dalley and Roiser 2012). Therefore, increased oculomotor impulsivity could be a consequence of altered serotoninergic transmission in depressed patients (Coppen 1967; see Yohn et al. 2017). The saccadic system is a high gain system constantly kept under strong inhibitory control in order to avoid unwanted eye movements (Missal and Keller 2002; Otero-Millan et al. 2018) and this delicate balance could be easily perturbed by reduced top-down inhibitory control (see review in Pouget et al. 2017).

Additionally, increased impulsivity could also be related to therapeutic drugs taken by subjects. Indeed, the RT-FP function of untreated subjects was similar to the one observed in low-impulsivity patients in general.

### Relationship between premature saccades and trait impulsivity

The absence of correlation between the percentage of premature saccades and *trait* impulsivity (BIS-11 total score) supports the hypothesis that impulsivity is a complex construct with different aspects and several different neurotransmitters involved (Evenden 1999; Dalley and Robbins 2017). Premature responses characterize the motor side of impulsivity but does not reflect higher order cognitive aspects of this phenomenon. When controls and patients were matched for impulsivity, temporal preparation was nevertheless reduced in the latter group. Therefore, impulsivity alone cannot explain observed results and depression by itself reduced temporal preparation. Figure 5 shows a schematic representation of the two opposing processes influencing saccadic latency during the FP. Oculomotor impulsivity reduces reaction time to the temporally proximal stimulus at the cost of more premature responses whereas depression reduces the influence of elapsed time on movement latency. Using the exquisite temporal sensitivity of the oculomotor system we have shown that the *implicit* processing of time is altered in MDD patients. This implicit processing of time probably involves early neuronal activity in the frontal cortex. It has been shown by Los and Heslenfeld (2005) that the effects of previous FP’s were paralleled by similar effects on the fronto-central Contingent Negative Variation (CNV). We suggest that this modulation of the CNV by sequence effects (or short-term temporal memory) should be reduced in MDD.

**Figure 5.**
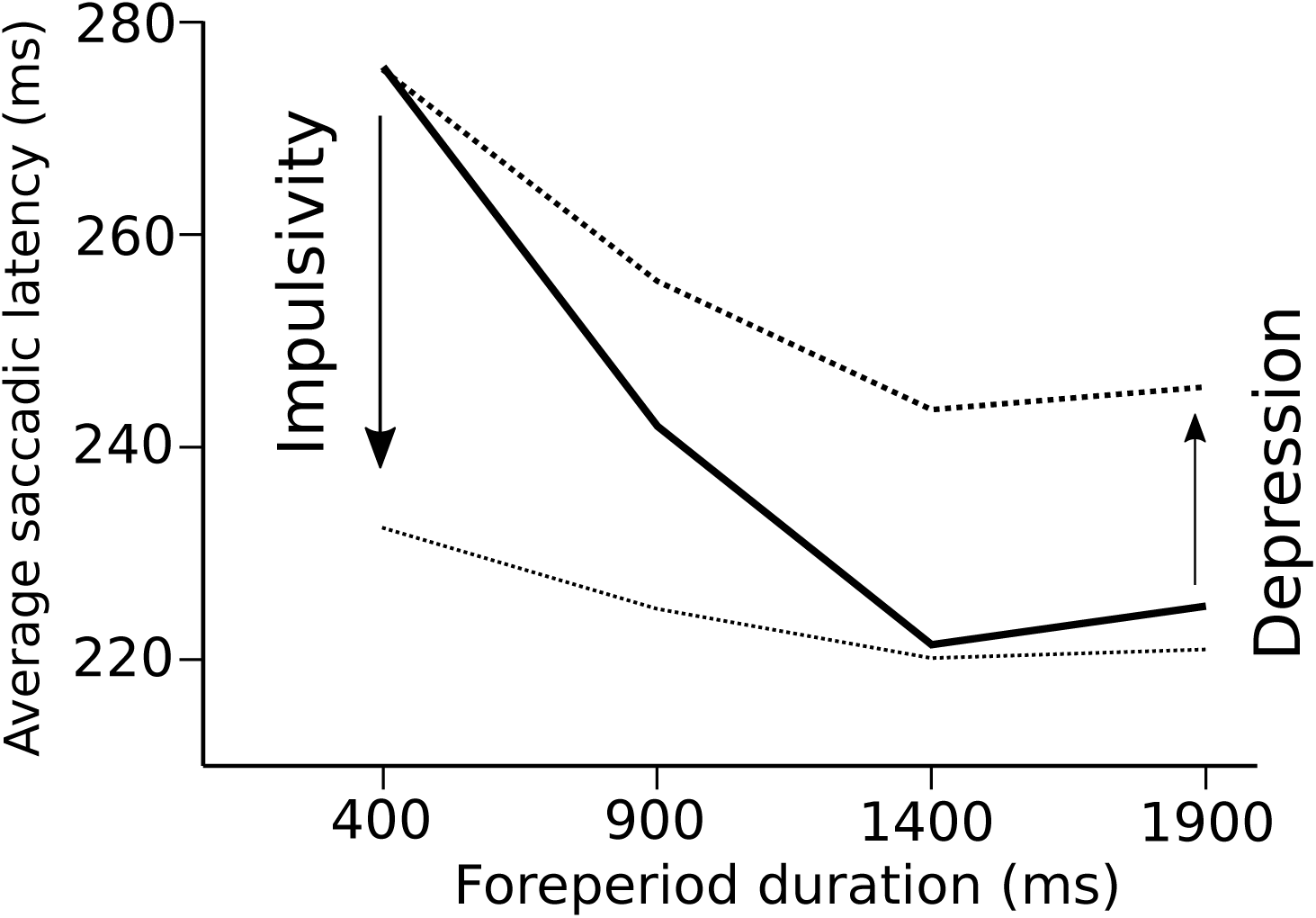
Schematic representation of the opposing influences of oculomotor impulsivity and depression on temporal preparation. Impulsivity decreases reaction times to targets appearing after a short FP (400 ms and 900 ms) but not to targets appearing after a long FP (1400 ms or 1900 ms). In contrast, depression does not alter saccadic latency after a short FP but increases latencies for all other FP durations.

### Implicit timing, time perception and awareness

Based on questionnaires, depressed patients report often a ‘slowing down’ of subjective time (Blewett, 1992; Ratcliff, 2012; see review in Droit-Volet, 2013). This desynchronization between the subjective experience of time and physical time seems to be a characteristic of depressive states without being systematically studied and compared with other cognitive functions. How temporal preparation, explicit timing and awareness of time relate to each other is unknown. Although temporal preparation and time awareness could be considered as very different phenomena they could rest on overlapping neurophysiological mechanisms. We suggest that alteration of these mechanisms could underlie the abnormal temporal organization of thought and behavior often reported in this disease.

## Acknowledgement

We would also like to express our gratitude to Professor Chi-Hung Juan for sharing equipment that greatly assisted the study. This manuscript has been released as a Pre-Print at bioRxiv.

## 14. Conflict of Interest

Declarations of interest: none

## 15. Funding sources

### Funding

This research was supported by Taiwan Ministry of Science and Technology grants 105-2632-H-038-001-MY3 and 104-2420-H-038-001-MY3, to TJL. 104-2410-H-038-012-MY2, 106-2410-H-038 -004 -MY2, to TYH. It was also supported by the Fund for Scientific Research (FRS-FNRS, Belgium) grant 31235458 (Research Credit) and an international mobility grant from Wallonie-Bruxelles International (WBI), to MM.

